# AC2P20 selectively kills *M. tuberculosis* at acidic pH by depleting free thiols

**DOI:** 10.1101/2021.03.16.435645

**Authors:** Shelby J. Dechow, Garry B. Coulson, Michael W. Wilson, Scott D. Larsen, Robert B. Abramovitch

## Abstract

*Mycobacterium tuberculosis* (Mtb) senses and adapts to host immune cues as part of its pathogenesis. One environmental cue sensed by Mtb is the acidic pH of its host niche in the macrophage phagosome. Disrupting the ability of Mtb to sense and adapt to acidic pH has the potential to reduce survival of Mtb in macrophages. Previously, a high throughput screen of a ∼220,000 compound small molecule library was conducted to discover chemical probes that inhibit Mtb growth at acidic pH. The screen discovered chemical probes that kill Mtb at pH 5.7 but are inactive at pH 7.0. In this study, AC2P20 was prioritized for continued study to test the hypothesis that it was targeting Mtb pathways associated with pH-driven adaptation. RNAseq transcriptional profiling studies showed AC2P20 modulates expression of genes associated with redox homeostasis. Gene enrichment analysis revealed that the AC2P20 transcriptional profile had significant overlap with a previously characterized pH-selective inhibitor, AC2P36. Like AC2P36, we show that AC2P20 kills Mtb by selectively depleting free thiols at acidic pH. Mass spectrometry studies show the formation of a disulfide bond between AC2P20 and reduced glutathione, supporting a mechanism where AC2P20 is able to deplete intracellular thiols and dysregulate redox homeostasis. The observation of two independent molecules targeting free thiols to kill Mtb at acidic pH further supports that Mtb has restricted redox homeostasis and sensitivity to thiol-oxidative stress at acidic pH.

## Introduction

Mtb pathogenesis is driven by its ability to exploit and adapt to the intracellular host environment. During pathogenesis, Mtb encounters a variety of stressors including nitrosative, oxidative, acidic pH, and hypoxic stress [1]. In response to these stresses, Mtb alters its physiology in order to survive the hostile macrophage environment and modulate expression of virulence genes critical for its pathogenicity. Acidic pH is an initial environmental cue that Mtb senses upon infection of the host macrophage [2, 3]. For survival within the resting macrophage, Mtb inhibits fusion of the phagosome and lysosome and resides in a mildly acidic phagosome (pH 6.4) [4]. Activation of the macrophage leads to phagosome acidification and Mtb resists this acid stress, maintaining a relatively neutral cytoplasmic pH, even at pH <5.0 [5–8]. In addition to expressing mechanisms to survive acid stress, Mtb also exhibits pH-and-carbon source dependent growth adaptations. Mtb will completely arrest its growth in minimal media buffered to pH 5.7 with glycerol as the sole carbon source [9]. During this growth arrest, Mtb exhibits carbon specificity, and will only arrest growth on glycolytic carbon sources (i.e. glucose and glycerol) [9]. However, when given specific carbon sources (i.e. phosphoenolpyruvate, pyruvate, acetate, oxaloacetate, and cholesterol), Mtb resuscitates its growth at pH 5.7 in minimal media, and thus, exhibits direct metabolic remodeling during pH stress [9]. Collectively, these studies show that in response to acidic pH, Mtb has multiple mechanisms in place whereby it alters its physiology for survival and virulence.

When Mtb is cultured at acidic pH or in macrophages, the bacterium has an imbalanced redox state with a more reduced cytoplasm [9, 10], a phenomenon referred to as reductive stress [3, 11]. It is hypothesized that acidic pH may cause redox imbalances due to adaptations of the electron transport chain that promote oxidative phosphorylation while maintaining cytoplasmic pH homeostasis [3]. These adaptations could lead to an accumulation of reduced co-factors such as NADH/NADPH. Implications for this type of reductive stress included altered Mtb metabolism, slowed growth, and non-replicating persistence. Fatty acid synthesis is thought to help mitigate reductive stress via the oxidation of NADPH and is supported by the induction of genes associated with lipid metabolism and anaplerosis at low pH [2,3,9,12]. One of these induced genes is WhiB3, a regulatory protein that senses Mtb’s intracellular redox state through its [4Fe-4S] cluster and acts to mitigate reductive stress [11,13,14]. WhiB3 is thought to counter this reductive stress via its role as a metabolic regulator, whereby it controls the anabolism of virulence lipids: poly- and diacyltrehalose (PAT/DAT), phthiocerol dimycocerosate (PDIM), and sulfolipids (SL-1) [14]. Production of these methyl-branched polar lipids requires NADPH; therefore, WhiB3 helps alleviate reductive stress by channeling excess reductants into fatty acid synthesis [14]. This results in the re-oxidation of reducing equivalents needed to maintain intracellular redox homeostasis. Changes in central metabolism, including the induction of anaplerotic pathways driven by isocitrate lyases (*icl*) and phosphoenolpyruvate carboxykinase (*pckA*) at acidic pH [15], and the dependence on carbon sources that feed the anaplerotic node [9], may also provide metabolic flexibility required to balance redox homeostasis at acidic pH.

Mechanisms important for pH adaptation (i.e. metabolism, cytoplasmic pH-homeostasis, and redox homeostasis) present an attractive source of novel targetable physiologies for drug discovery. pH homeostasis can be targeted by compounds like the benzoxazinone, BO43, which inhibits the serine protease MarP, resulting in the disruption of intrabacterial pH homeostasis [16]. Additionally, ionophores have also been discovered to kill Mtb at acidic pH [17, 18]. Respiration has been shown to be important for maintaining pH-homeostasis [19, 20]. Compounds targeting respiration include Bedaquiline (BDQ), a F_1_F_o_-ATP-synthase inhibitor, and the small molecule, C10. BDQ has been shown to act as an ionophore and disrupt the Mtb transmembrane pH gradient [21], while C10 exhibits enhanced Mtb killing at acid stress [22]. Thiol-redox homeostasis also has implications as a targetable pH-dependent physiology. Auranofin depletes free thiols by targeting an essential thioredoxin reductase (TrxB2) [23, 24]. Together, these results demonstrate the druggability of physiologies important for acidic pH-dependent adaptation.

PhoPR, a two-component regulatory system (TCS), is important for regulating Mtb virulence and intracellular survival [25,12,26]. Additionally, signaling from PhoPR has been shown to play an important role in pH adaptation [2,27,9]. Our lab previously conducted a reporter based, whole cell high-throughput screen (HTS) of > 220,000 small molecules for inhibitors of PhoPR signaling at acidic pH [28, 29]. Compound activity was assessed in rich media buffered to pH 5.7 using a pH-inducible Mtb fluorescent reporter strain to identify either direct inhibitors of the PhoPR regulon or pH-selective inhibitors of Mtb growth. This screen successfully identified inhibitors of *phoPR*-dependent signaling, including the carbonic anhydrase (CA) inhibitor, ethoxzolamide (ETZ) [28]. This screen also identified compounds that selectively kill Mtb at pH 5.7 but not pH 7.0 and do so independently of PhoPR. One of these compounds, called AC2P36 (5-chloro-N-(3-chloro-4-methoxyphenyl)-2-methylsulfonylpyrimidine-4-carboxamide) [29], functions by directly depleting intracellular Mtb thiol pools, by forming covalent adducts with free thiols. Depletion of free thiols, interferes with redox buffering pathways and induces formation of cytoplasmic reactive oxygen species (ROS) at acidic pH, thus sensitizing Mtb to thiol-oxidative stress [29]. AC2P36 also selectively kills Mtb and potentiates the activity of TB drugs: isoniazid, clofazimine, and diamide. We hypothesize that reductive stress at acidic pH selectively sensitizes Mtb to thiol targeting activity of AC2P36. These results indicate that free thiols are a pH-selective target, and that Mtb sensitivity to killing is enhanced under thiol oxidative stress.

In this study, we report on a new chemical probe isolated from a prior screen, AC2P20 (N-1,3-benzothiazol-2-yl-2-[(4,6-dioxo-5-phenyl-1,4,5,6-tetrahydropyrimidin-2-yl)thio]acetamide) (Figure 1A), that selectively kills Mtb at acidic pH. AC2P20 was identified as a *phoPR*-independent, pH-selective inhibitor of Mtb growth. Through transcriptional profiling we observed that genes modulated by AC2P20 treatment significantly overlap with genes modulated by AC2P36 treatment. Although both compounds are structurally distinct, like AC2P36, AC2P20 also exhibits killing of Mtb at pH 5.7, disrupts thiol homeostasis by depleting intracellular free thiol pools, and increases reactive oxygen ROS production. Thus, AC2P20 is a second structurally unique pH-selective chemical probe that exhibits thiol-depletion as a mechanism-of-action for killing at acidic pH. This finding further reinforces the vulnerability of Mtb to perturbations of redox homeostasis at acidic pH.

**Figure 1.**
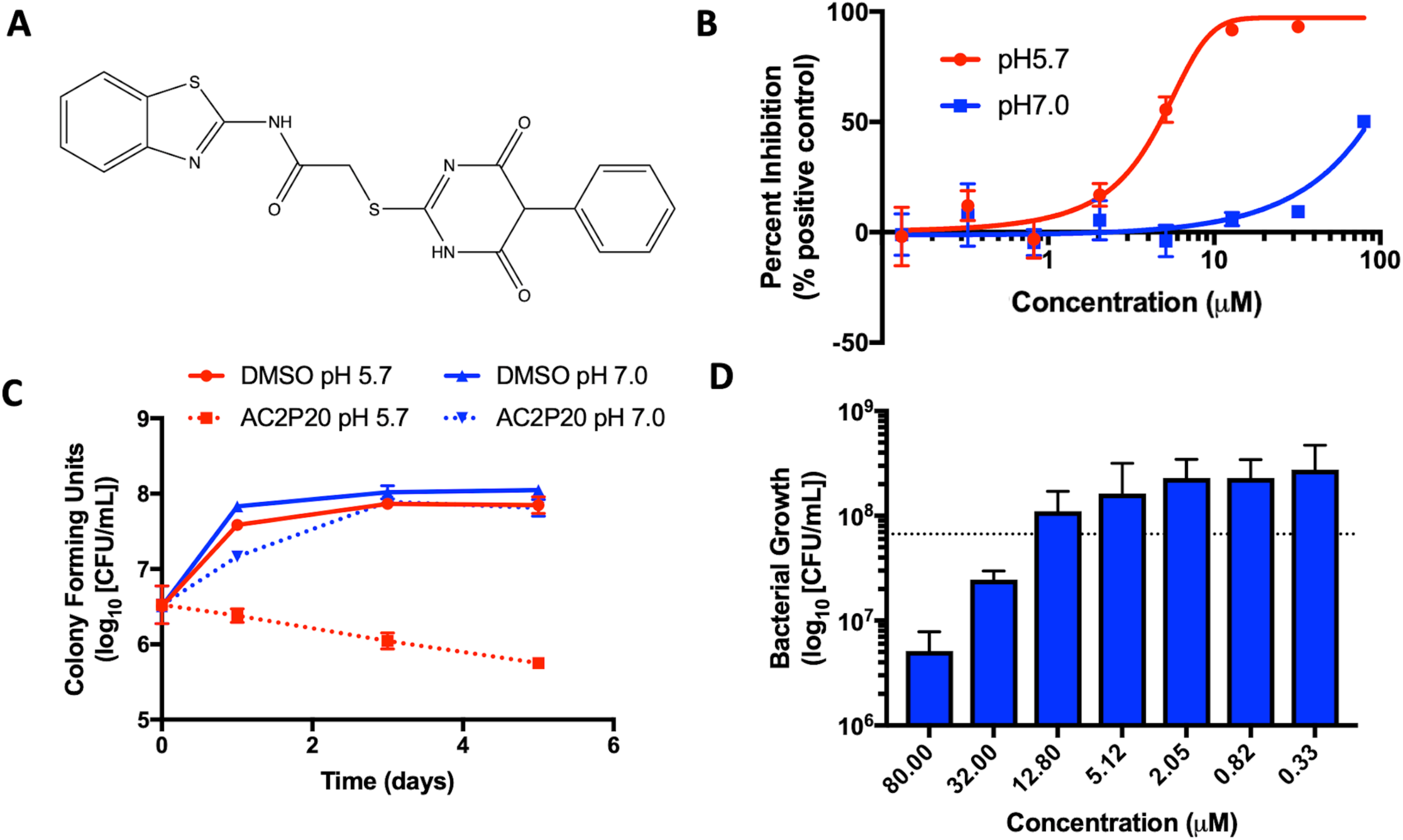
AC2P20 inhibits Mtb growth in a pH-dependent manner. A) The chemical structure of AC2P20 ((N-1,3-benzothiazol-2-yl-2-[(4,6-dioxo-5-phenyl-1,4,5,6-tetrahydropyrimidin-2-yl)thio]acetamide) B) Mtb growth is inhibited in a dose-dependent manner when treated with AC2P20 at pH 5.7 and exhibits an EC_50_ of 4.3 μM following six days of treatment. Treatment with AC2P20 at pH 7.0 Mtb requires concentrations >60 μM to see growth inhibitory effects. C) Mtb treated with 20 μM of AC2P20 and grown in buffered 7H9 media (pH 5.7) for 5 days shows time-dependent killing as indicated by ∼100-fold reduction in viability compared to the DMSO control. Time-dependent killing is not observed in neutral conditions. D) Mtb was treated with a dose-response of AC2P20 at pH 5.7 for 7 days, then assessed for dose-dependent killing by plating for colony-forming units (CFUs). The dotted line indicates the CFUs plated on Day 0.

## Experimental

### Bacterial Strains and Growth Conditions

*M. tuberculosis* strains Erdman and CDC1551 and *M. smegmatis* strain mc^2^155 (expressing GFP from a replicating plasmid) were used in all experiments unless specified. Mtb was cultured in Middlebrook 7H9 media enriched with 10% oleic acid-albumin-dextrose-catalase (OADC), 0.05% Tween-80, and glycerol. Cultures were maintained in vented T-25 culture flasks and grown at 37 °C and 5% CO_2_. To maintain a specific pH, 7H9 media was strongly buffered to pH 7.0 with 100 mM 3-(N-morpholino)propanesulfonic acid (MOPS) or pH 5.7 with 100 mM 2-(N-morpholino)ethanesulfonic acid (MES). Mtb was grown to mid-late log phase (OD_600_ 0.5-1.0) before exposure to buffered 7H9 for use in experiments detailed below. *M. smegmatis* cultures were grown in identical 7H9 media conditions at a starting OD_600_ 0.05 at 37°C in a shaking incubator (200 rpm).

### Selection for AC2P20 resistant mutants

Mtb CDC1551 and Mtb Erdman strains were grown to an OD_600_ of 0.6-1.0, spun down, and resuspended in 7H9 media buffered to pH 5.7. Mtb cells were plated at 10^9^ cells/mL on 7H10 agar media buffered to pH 5.7 and supplemented with 10 µM, 20µM or 40 µM AC2P20. Plates were incubated at 37°C for over 12 weeks without any significant isolated colonies appearing. This experiment was performed three times with similar results.

### Transcriptional profiling and data analysis

Mtb cultures were grown at 37°C and 5% CO2 in standing T-25 culture flasks to an OD_600_ of 0.5 in 8 mL of 7H9 buffered media. Treatment conditions examined include i) 20µM AC2P20 at pH 5.7 and ii) an equivalent volume of DMSO at pH 5.7 as the baseline control. Each culture was incubated for 24 hours and treatment conditions were conducted in two biological replicates. Following incubation, total bacterial RNA was extracted as previously described [2, 9] and sequencing data was analyzed using SPARTA (ver. 1.0) [30]. Genes identified filtered based on log_2_CPM < 5 and log_2_FC < 1. A Chi-Square analysis with Yates Correction was conducted to test the statistical relationship between gene overlap with the AC2P36 transcriptional profile as described by Coulson *et al.* [29]. The RNAseq data has been deposited at the GEO database (Accession # GSE151884).

### Half-maximal effective concentration (EC_50_) determination and spectrum of activity in other mycobacteria

Mtb cultures were incubated in buffered 7H9 media (pH 5.7 or pH 7.0) at a starting OD_600_ of 0.2, with 200 uL aliquoted into 96-well microtiter assay plates (CoStar #3603). Cultures were treated with a 2.5-fold dose-response of AC2P20 (80 µM-0.13 µM) and incubated standing for 6 days at 37 °C and 5% CO_2_, with bacterial growth assessed by optical density (OD_600_). Cultures treated with an equivalent volume of DMSO or 0.3 µM Rifampin were used as negative and positive controls, respectively. Each condition was performed in duplicate and representative of three individual experiments. EC_50_ values were determined using GraphPad Prism software (ver. 7.0). AC2P20 activity against *M. smegmatis* was also performed in 96-well assay plates in 7H9 buffered media (pH 7.0 or 5.7). *M. smegmatis* cultures were seeded at a starting OD_600_ of 0.05 with 200 uL aliquoted into each well. An 8-point 2.5-fold dilution series starting at 80 µM was conducted and cultures were incubated for 3 days with shaking (100 rpm). Plates were read for GFP fluorescence.

### Mycobactericidal activity of AC2P20

Mtb was initially cultured in 7H9 media (pH 5.7 or 7.0) at a starting OD_600_ of 0.2 in 96-well assay plates. Cultures were treated with a 2.5-fold dose-response of AC2P20 (80 µM-0.33 µM). An equivalent volume of DMSO was included as a control. Each treatment condition was conducted in triplicate and incubated for 7 days. Following incubation, treated wells were serially diluted in 1X Phosphate-Buffered Saline (PBS) and plated for colony forming units (CFUs) on 7H10 agar plates supplemented with 10% OADC and glycerol. Bactericidal activity was determined by comparing CFUs from the initial inoculum to CFUs following treatment.

### Cytoplasmic pH-homeostasis

Mtb washed with PBS (pH 7.0) was labelled with Cell Tracker 5’-chloromethylfuoroscein diacetate (CMFDA) and analyzed using methods previously described [31]. Mtb treated with AC2P20 in PBS (pH 5.7) was assayed for cytoplasmic pH changes over the course of 24-hours. Excitation ratio results were converted to pH via a standard curve generated using Nigericin-treated Mtb in buffers of known pH. Treated Mtb results were then compared to the DMSO and Nigericin negative and positive controls, respectively.

### Measurement of endogenous reactive oxygen species

CellROX Green fluorescent dye (Invitrogen) was used to detect accumulation of endogenous reactive oxygen species (ROS) in Mtb as previously reported [29, 32]. Mtb grown to mid-late log phase was pelleted and re-suspended at a starting OD_600_ of 0.5 in 5 mL of buffered 7H9 media (pH 5.7 or 7.0) lacking catalase. Cultures prepared in duplicate were treated with two separate concentrations of AC2P20 (2 µM and 20 µM) and incubated for 24 hours at 37 °C. Following treatment, cultures were incubated with 5 mM CellROX Green (Thermo Fisher) for 1 hour at 37 °C and then washed twice with 1X PBS + 0.05% Tween80. Washed cells were resuspended in 0.6 mL 1X PBS and aliquoted into triplicate wells in 96-well microtiter plates. Wells were measured for fluorescence and optical density, with florescence being subsequently normalized to cell growth for ROS analysis. AC2P36 (2 µM and 20 µM) and equivalent volumes of DMSO served as positive and baseline controls, respectively.

### Detecting intracellular free thiol pools

Mtb grown in 7H9 OAD media lacking catalase was inoculated at a starting OD_600_ of 0.25 in 8 mL of buffered 7H9 OAD media (pH 5.7 or 7.0) also lacking catalase. Cultures were prepared in duplicate and treated with either DMSO, 2 µM AC2P20, 20 µM AC2P20, 20 µM AC2P36, or 20 µM auranofin. Treated cultures were incubated for 24 hours at 37 °C, normalized by OD_600_, and washed twice in 1X PBS supplemented with 0.05% Tyloxapol. Cells were resuspended in 0.75 mL of thiol assay buffer (100 mM potassium phosphate pH 7.4, and 1 mM EDTA) and lysed by bead beating for 2 minutes at room temperature. Supernatants were removed and saved for analysis using the Cayman Thiol Detection Assay Kit (Caymen Chemical) as previously described [29]. Thiol concentrations were measured in (nM) against a glutathione standard.

### Mass Spectrometry

Mass spectrometry was used to detect the formation of AC2P20 adducts. Aqueous solutions of 80 µM AC2P20 were prepared separately and incubated with either reduced glutathione (100 µM), N-acetylcysteine (100 µM), or hydrogen peroxide (100 µM) for 1 hour at room temperature in Tris-HCl buffer (pH 5.7, 7.0, or 8.5). Samples were analyzed using the Waters Xevo G2-XS QTof mass spectrometer (Milford, MA, USA) in both positive and negative electrospray ionization (ESI) modes. Samples were run with the following ion source parameters: capillary voltage, 2 kV; sampling cone, 35 V; source temperature, 100°C; desolvation temperature, 350°C; cone gas flow, 25 L/h; desolvation gas flow, 600 L/h. Ultra-performance liquid chromatography (UPLC), using water and acetonitrile as solvents, was carried out for the chromatographic separation of compounds. The LC parameters were as follows: flow rate, 0.2 mL/min; water/acetonitrile solvent gradient, 50/50 for 2 min. Mass analysis was performed at <1,500 Da. This experiment was repeated twice in duplicate with similar results seen at both positive and negative ESI.

## Results

### AC2P20 exhibits pH-dependent growth inhibition of M. tuberculosis

Two high throughput screens (HTS) using Mtb fluorescent reporters were conducted in order to detect inhibitors of two separate Mtb two-component regulatory systems (TCS): DosRST and PhoPR [28,29,33,34]. A chemical library of >220,000 small molecules was previously screened, with compound hits being defined as those that inhibited reporter fluorescence or Mtb growth. These compounds were further classified as TCS target inhibitors or growth inhibitors. The screens only differed in the reporter strain used and the pH of the medium, which was neutral or acidic in the DosRST and PhoPR inhibitor screens, respectively. Comparing growth inhibiting hits from these two screens identified a subset of compounds that selectively inhibited Mtb growth at acidic pH independent of PhoPR signaling. These compounds were classified as pH-selective growth inhibitors if they exhibited >50% growth inhibition at acidic pH and < 10% inhibition at neutral pH. AC2P20 (N-1,3-benzothiazol-2-yl-2-[(4,6-dioxo-5-phenyl-1,4,5,6-tetrahydropyrimidin-2-yl)thio]acetamide) (Figure 1A) exhibited >5-fold selectivity at acidic pH and was characterized as one of these pH-selective inhibitors of Mtb growth. The pH-dependent activity of AC2P20 was confirmed by determining its half-maximal effective concentration (EC_50_). Mtb treated with an 8-point dose-response of AC2P20 for six days at pH 5.7 results in dose-dependent growth inhibition with an EC_50_ of 4.3 μM, however, has a >10-fold higher EC_50_ of ∼60 μM at pH 7.0 (Figure 1B). AC2P20 also exhibits mycobacterial selectivity for Mtb compared to *M. smegmatis*, which has an EC_50_ > 80 μM at acidic pH and does not exhibit growth inhibitory activity at neutral pH (Figure S1A).

Time-dependent and concentration-dependent killing assays were conducted to define whether AC2P20 is bactericidal or bacteriostatic. Mtb treated with 20 μM AC2P20 exhibits pH-selective inhibition of Mtb growth in acidic conditions and results in approximately 2-log fold reduction in CFUs over 5 days (Figure 1C). In contrast, DMSO controls and AC2P20 treatment in neutral conditions have no impact on growth. The concentration-dependent killing assay shows that AC2P20 is bactericidal at ∼32 μM and bacteriostatic at 12 μM (Figure 1D). Cytoplasmic pH was measured to determine whether AC2P20 functions as an ionophore. Treatment with AC2P20 does not modulate the cytoplasmic pH of Mtb compared to the nigericin positive control (Figure S1B). Together, these data show that AC2P20 activity is pH-dependent, bactericidal, and does not alter Mtb cytoplasmic pH homeostasis.

### AC2P20 induces a thiol oxidative stress response similar to AC2P36

To isolate resistant mutants and thereby find potential targets for AC2P20, 10^9^ Mtb cells were plated on 7H10 agar media buffered to pH 5.7 containing 10 µM, 20 µM or 40 µM AC2P20. Despite several weeks of incubation each time at 37°C, no spontaneous mutants were isolated from multiple rounds of screening for resistant mutants to AC2P20. Following our resistance screening attempts, transcriptional profiling was conducted to define Mtb physiologies targeted following AC2P20 treatment. Mtb CDC1551 cultures were prepared in rich media (pH 5.7) and treated with 20µM AC2P20 or DMSO control for 24 hours. Mtb treated with AC2P20 caused induction of 156 genes (>2-fold, q < 0.05) and repression of 81 genes (>2-fold, q < 0.05) (Figure 2A, Table S1). Using MycoBrowser [35] to classify gene function, we found that the functional pathway most induced by AC2P20 (excluding conserved hypotheticals) was intermediary metabolism and respiration (Figure 2B, Table S1A, Table S1B). Differentially induced genes included genes involved in sulphur metabolism (*cysT, sirA, mec*), transcriptional regulation of the stress response (*sigH, sigB, rshA*), and redox homeostasis (*katG, trxB1, trxC*) (Figure 2C, Table S1B). Notably, differentially regulated genes from AC2P20 treated cells overlapped with differential gene expression profiles previously characterized for the pH-selective Mtb growth inhibitor, AC2P36 [29]. Gene enrichment analysis showed a statistically significant overlap between groups AC2P20 and AC2P36 differentially expressed genes (p<0.0001) (Figure 2C). Based on RNAseq data and the gene enrichment analysis, both AC2P36 and AC2P20 exhibit a transcriptional profile indicative of redox and thiol-oxidative stress. For example, both transcriptomes show induction of the alternative sigma factor *sigH* regulon which plays a central role in regulating thiol-oxidative stress during Mtb pathogenesis [36–38]. *sigH* is responsible for regulating genes involved in thiol metabolism including thioredoxin (*trxC*), thioredoxin reductases (*trxB1, trxB2*), and cysteine biosynthesis and sulfate transport (*cysO*, *cysM, cysA, cysW, cysT*). Additionally, *sigH*-regulated *moeZ* is induced in both transcriptomes, which is involved in sulfation of enzymes and plays a dual role in molybdopterin biosynthesis and *cysO* activation[39]. While the *sigH* regulon exhibits a direct response to thiol-oxidative stress, it is also highly induced under oxidative stress conditions[36]. In addition, non-*sigH* regulated oxidative stress responsive genes include *katG* (catalase-peroxidase), *thiX* (a thioredoxin), and *furA* (transcriptional regulator), which are upregulated in both AC2P20 and AC2P36. Interestingly, *Rv0560c*, a methyltransferase, is the most upregulated gene in Mtb treated with AC2P20, AC2P36, or C10 [22, 29]. *Rv0560c* is induced in mutants resistant to a cyano-substituted fused pyrido-benzimidazole, known as compound 14, and provides resistance by methylating and inactivating compound 14 [40]. *Rv0560c* is not directly upregulated by the *sigH* regulon or oxidative stress, but rather by salicylate [41], and may be involved in the synthesis of redox cycling agents [42–44]. Therefore, induction of thiol-homeostasis metabolism genes and *katG* in response to AC2P20 treatment suggests an increased need for the generation of low molecular weight thiols, which are important for detoxification of toxic reactive oxygen species (ROS) and maintaining redox homeostasis in Mtb.

**Figure 2.**
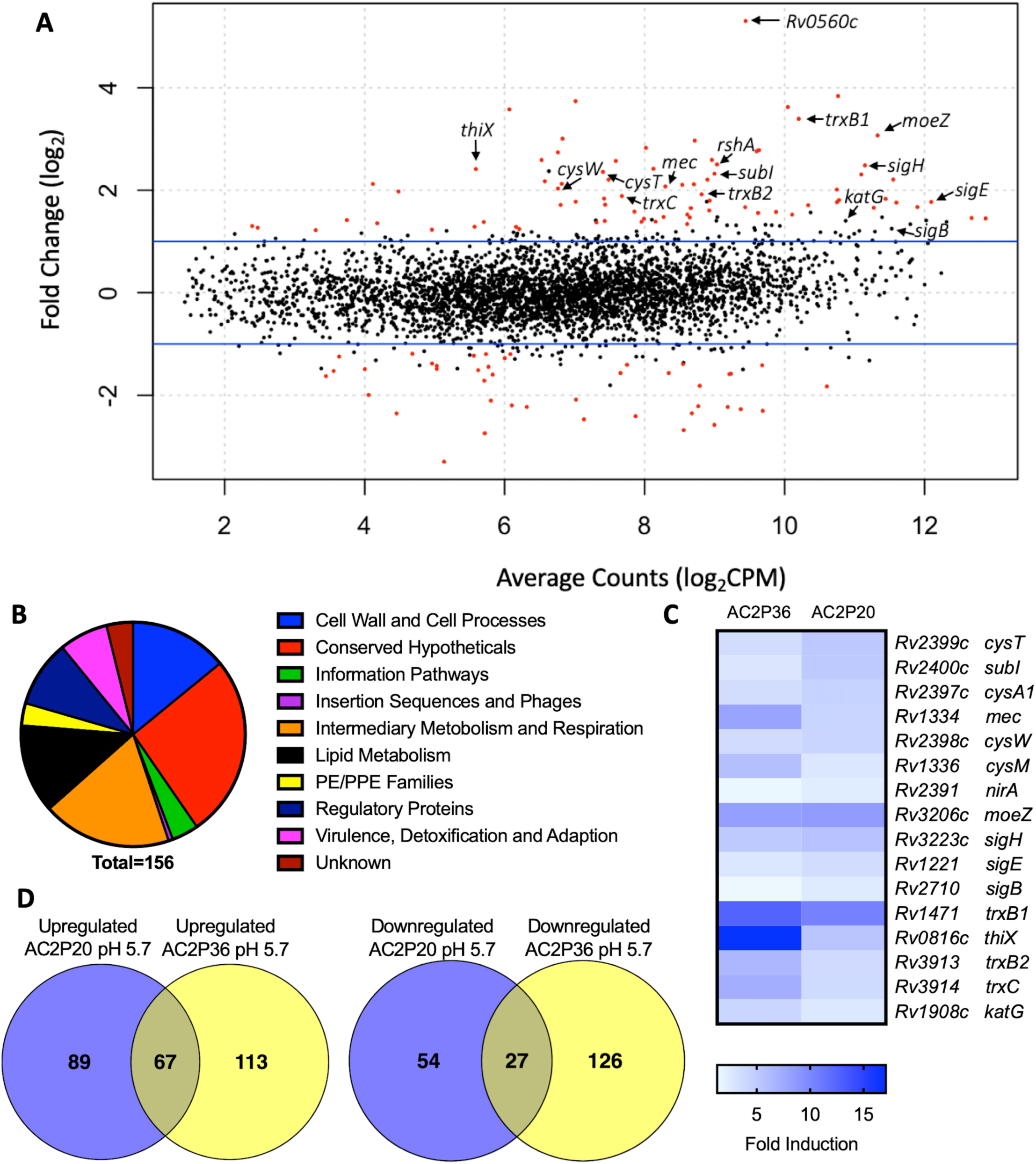
AC2P20 treatment promotes a thiol-oxidative and redox stress response. A) Mtb differential gene expression data after being treated for 24 hours with 20 µM AC2P20 at pH 5.7. Genes indicated include those involved in sulfur metabolism, transcriptional regulation, and redox homeostasis. Statistically significant genes (q < 0.05) are highlighted in red. B) A pie chart depicting the functional classification breakdown of significantly induced genes (>2-fold, q < 0.05) following the analysis of AC2P20-treated Mtb RNA-seq profile. C) Heatmap comparing 16 upregulated genes (between AC2P20 and AC2P36 at pH 5.7 that are involved in sulfur metabolism, transcriptional regulation, and redox homeostasis. Genes were annotated with the H37Rv genome. D) Venn diagrams comparing upregulated and downregulated gene overlap (>2-fold, q < 0.05) between AC2P20-treated and AC2P36-treated Mtb ^29^.

Despite significant overlap between the AC2P20 and AC2P36-treated regulons, there are pathways that are distinctly different in the transcriptional profiling comparisons. Classification of gene function for the 180 AC2P36-induced genes (>2-fold, q < 0.05) showed that the functional category most induced (excluding conserved hypotheticals) was intermediary metabolism and respiration, the same as AC2P20. However, major differences were noted between categories of both induced gene sets for AC2P20 and AC2P36. For example, induction of lipid metabolism genes comprised roughly 3.33% of the total genes induced by AC2P36 compared to 12.82% for AC2P20 (Figure 2B, Figure S2A). Noticeably, AC2P20 appeared to upregulate several mycolic acid biosynthesis pathway and operon genes (*fas, acpM, kasA, accD6*) (Figure S2B). In contrast, these genes were repressed following AC2P36 treatment. Other lipid metabolism genes not observed in AC2P20 transcriptional data, but actively repressed by AC2P36 include *scoA/B*, *accD1*, *Rv3087*, and *fadE35* [29]. Additionally, transcriptional profiling showed that methylcitrate synthase and methylcitrate dehydratase genes (*prpC* and *prpD*, respectively) were oppositely modulated in both regulons; AC2P20 repressed *prpC/D* expression while their expression was induced by AC2P36 (Figure S2B). Other functional categories that saw large quantitative changes between both transcriptional profiles include cell wall and cell wall processes and virulence, detoxification and adaptation. Fewer cell wall and cell wall processes genes were induced by AC2P36 compared to AC2P20, while the number of virulence, detoxification and adaptation functional genes were increased following AC2P36 treatment (Figure S2A). The transcriptional differences observed between both regulons demonstrates that despite the shared similarities in regulation of thiol-redox homeostasis and regulatory genes, distinct differences exist between how pathways are modulated following AC2P20 and AC2P36, with lipid metabolism being most notable.

### AC2P20 forms an adduct with the low molecular weight thiol, GSH

Although AC2P36 and AC2P20 have distinct structures, both compounds contain a similar thiol-containing pyrimidine group. In AC2P36, it is thought that the methylsulfone moiety acts as an electron-withdrawing group which allows a thiolate anion to undergo nucleophilic attack on the C-2 carbon of the pyrimidine ring in order to release methanesulfinic acid or methanesulfinate (Figure S1C) [29]. This interaction is thought to result in the formation of a sulfide bond and depletion of available free thiols. Indeed, heteroaromatic sulfones have been recently described as tunable agents for cysteine-reactive profiling [45, 46]. Based on these observations with AC2P36, and the noted similarity with the thiol-containing pyrimidine group, we hypothesized that AC2P20 may have a similar mechanism of action and undergo covalent modification of free thiols. To test this hypothesis, 80µM AC2P20 was incubated with 100µM reduced glutathione (GSH) for one hour in basic, neutral, and acidic conditions and analyzed via mass spectrometry. Incubation of AC2P20 with GSH resulted in the formation of an adduct at pH 5.7 with an exact molecular weight of ∼529 Da (Figure 3A, Figure 3C, Figure S5). There is also adduct formation in neutral and basic conditions (Figure S3A, Figure S3B) although with lower peak intensity. AC2P20 incubated with DMSO does not appear to fragment in the absence of GSH in any of these conditions. (Figure 3B, Figure S3C, Figure S3D). In the positive ESI mode (Figure 3C), a neutral fragment of 129 Da is lost from the adduct with a peak seen at ∼401 Da, consistent with a loss of the glutamate fragment from GSH [47]. Fragmentation of AC2P20 is also observed when incubated with GSH at pH 5.7, with peaks at ∼222 Da, ∼206 Da, ∼194 Da, and ∼178 Da aligning with possible fragments of the pyrimidine group of AC2P20 (Figure S5). The peak observed at ∼391 Da is a mass spectrometry plasticizer and common contaminant that can be used for mass calibration [48]. We also looked at N-acetylcysteine (NAC), a derivative of GSH, and its ability to form an adduct with AC2P20. A peak was observed at ∼384 Da, aligning with the formation of an AC2P20-NAC adduct (Figures S4A, Figure S5). Interestingly, higher peak intensities of these adducts were observed at neutral and basic conditions (Figure S4B, Figure S4C). This is possibly due to NAC having a pKa ∼9.5, and therefore favoring the adduct reaction with AC2P20 under these conditions. Together, these findings support that AC2P20 reacts with low molecular weight thiols and thiol groups. Additionally, we looked at whether AC2P20 still form an adduct with GSH in the presence of the oxidant, H_2_O_2_. It was thought that H_2_O_2_ may cause the formation of intermediate sulfenic acid and oxidize GSH, resulting in the formation of glutathione (GSSG) [49]. After incubating AC2P20 with both GSH and H_2_O_2_, we still observed disulfide bond formation between AC2P20 and GSH, indicating that GSSG is probably not being produced (Figure S4D). These results suggest that AC2P20 is capable of forming a disulfide bond with low molecular weight thiols.

**Figure 3.**
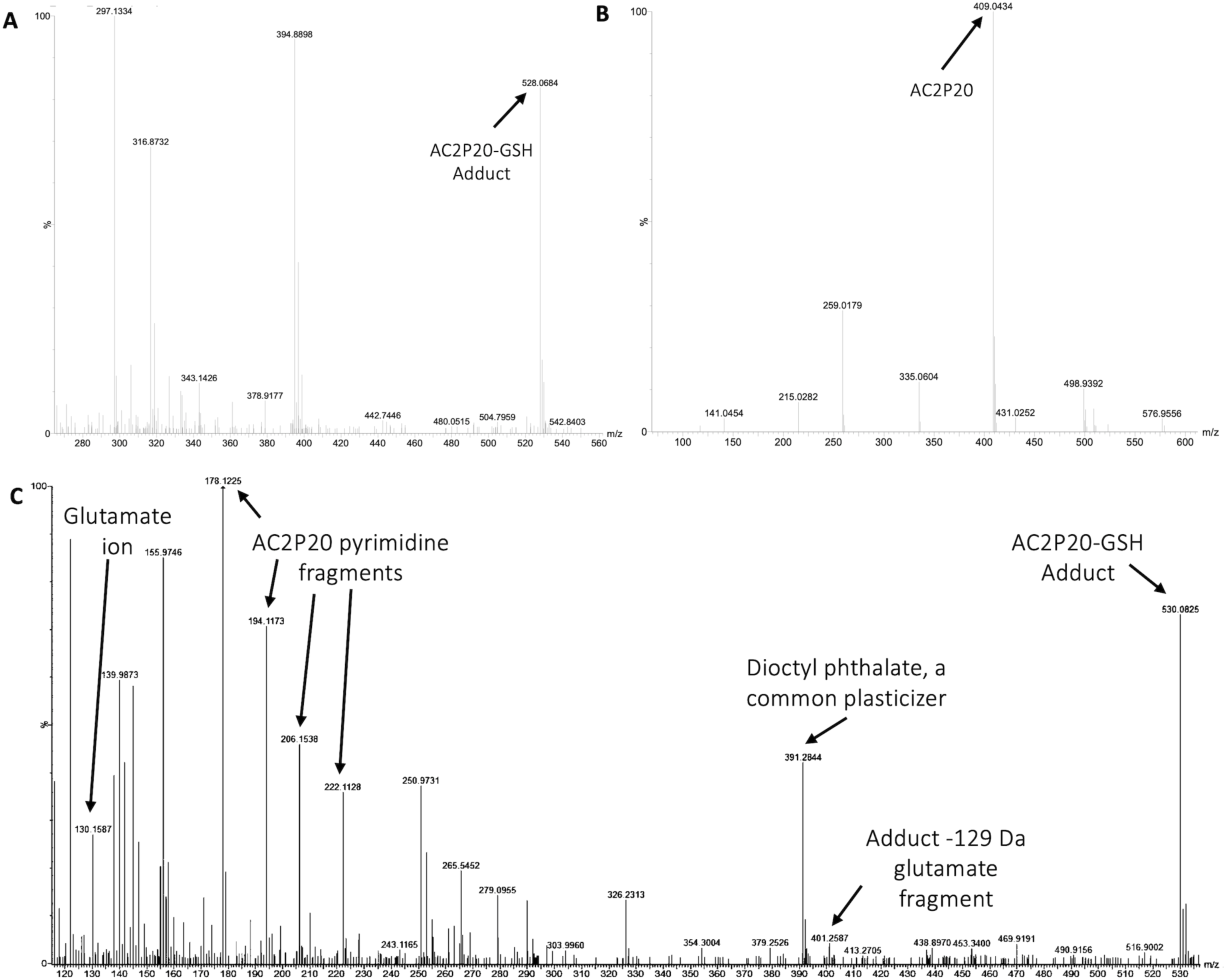
AC2P20 forms adducts with free thiols at acidic pH. A) AC2P20 was incubated in Tris-HCl buffer, pH 5.7 with reduced glutathione (GSH) for one hour. An AC2P20-GSH adduct (∼528 Da) was confirmed via mass spectrometry. Samples were run in duplicate and observed in negative ESI mode. B) In the absence of GSH, AC2P20 incubated with DMSO does not fragment at pH 5.7. Only the parent molecule is observable at a molecular weight of ∼409 Da. Samples were run in duplicate and observed in negative ESI mode. C) AC2P20-GSH adduct formation at pH 5.7 (∼530 Da) was also observed in positive ESI mode, as well as adduct loss of the glutamate fragment (∼401 Da) and subsequent fragmentation of the AC2P20 molecule and its pyrimidine fragments. Samples were run in duplicate.

### AC2P20 depletes free thiols and causes an accumulation in ROS in Mtb at acidic pH

Given that an adduct is able to form between AC2P20 and GSH, we sought to test the ability of AC2P20 to deplete free thiols in Mtb. For this assay, Mtb was treated with AC2P20 (2 μM and 20 μM) in both acidic and neutral conditions for 24 hours. Auranofin (20 μM) was used as a positive control because it inhibits Mtb’s thioredoxin reductase, TrxB2, thereby disrupting thiol- and redox-homeostasis[23]. AC2P36 (20 μM) was also included in the assay to compare thiol depleting activities of both compounds. Following AC2P20 treatment, a statistically significant reduction in free thiol concentrations was observed intracellularly in Mtb at pH 5.7 where free thiols are reduced by ∼2.8-fold to ∼133nM compared to the DMSO vehicle control at ∼380 nM (Figure 4A). As expected, we also see free thiol depletion in Mtb following treatment with both positive controls, supporting the observation seen with AC2P20. In contrast to Auranofin, AC2P20 treatment at neutral pH does not exhibit any statistically significant reduction in free thiols, supporting the pH-selective activity of this compound. Interestingly, AC2P36 does exhibit some activity in neutral conditions. This is possibly due to AC2P36 still exhibiting some growth inhibitory activity at neutral pH at ∼30 μM, whereas AC2P20 requires much higher concentrations (∼60 μM) to see a similar inhibitory effect.

**Figure 4.**
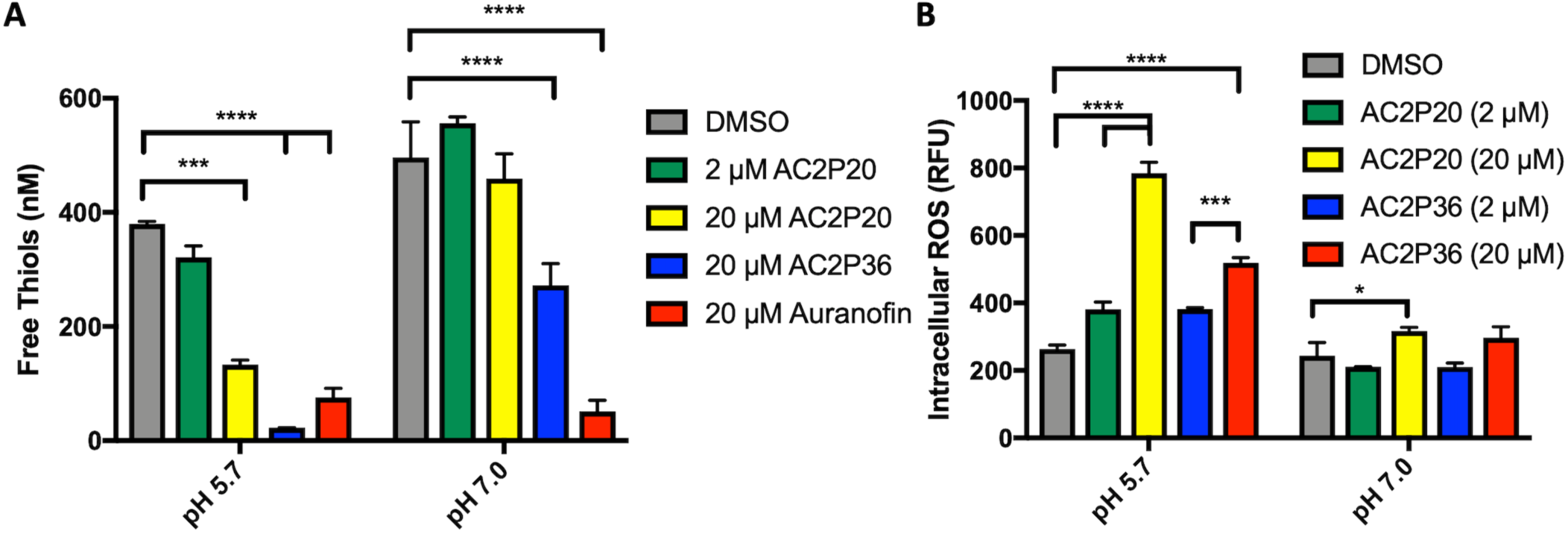
AC2P20 depletes free thiols and induces intracellular ROS accumulation. **A) Treatment of Mtb with AC2P20 leads to a pH-dependent decrease in free thiols.** Free thiol depletion is observed at pH 5.7 with AC2P20 treatment. AC2P36 is a pH-dependent chemical probe known to deplete free thiol pools and serves as a positive control. Statistical significance was calculated using a two-way ANOVA (*p<0.05). **B) ROS accumulate under AC2P20 treatment at acidic conditions.** Mtb treatment with AC2P20 leads to a pH-dependent increase in intracellular reactive oxygen species (ROS). ROS was detected using a final concentration of 5 µm fluorescent dye, CellROX Green, and normalized to an OD_595_. DMSO was used as a control. Statistical significance was calculated using a one-way ANOVA (*p<0.05)

Depletion of total free thiols will result in disrupted redox homeostasis and therefore may result in enhanced ROS accumulation. To test this hypothesis, we conducted an assay measuring intracellular ROS production in Mtb. Mtb was incubated with 2 μM and 20 μM AC2P20 for 24 hours, treated with CellROX fluorescent dye for 1 hour, and then assayed for relative fluorescence and optical density. AC2P36 (2 μM and 20 μM) was included as the positive control, because it has previously been shown to accumulate intracellular ROS following treatment. At acidic pH, 20 μM AC2P20 exhibits ∼3-fold increase in intracellular ROS production compared to DMSO (Figure 4B). AC2P20 (20 μM) also increases ROS accumulation ∼3-fold greater in acidic conditions compared to neutral pH, where there is little ROS accumulation compared to DMSO. AC2P36 (20 μM) also increases ROS production ∼2-fold at pH 5.7, which is consistent with previous observations. These data support a mechanism whereby enhanced ROS accumulation can be driven by pH stress and is further exacerbated by AC2P20 treatment.

## Discussion

Based on the chemical structure of AC2P20 and the adduct it forms with GSH at pH 5.7, we propose a reaction model where the benzothioazole-mercaptoacetamide group covalently modifies free thiols, forming stable adducts. Shown here is a potential mechanism for the generation of adducts observed by mass spectrometry (Figure 3A, Figure 3C). Disulfide bond formation between GSH (307.32 Da) and the free benzothioazole-mercaptoacetamide group (223.29 Da) results in a molecule mass of 529 Da, which can be observed in both positive and negative ESI modes (Figure 5A, Figure S5). Loss of the neutral glutamate fragment from the AC2P20-GSH adduct results in a peak at 401 Da (ESI+). We suspect AC2P20 may be undergoing hydrolysis, however, we do not observe the phenyl-dioxopyrimidine fragment (204 Da). We do observe a fragmented phenyl-dioxopyrimidine group at 178 Da which may be due to the sample’s molecules breaking into charged fragments during mass spectrometry. The absence of a 204 Da fragment may also suggest that adduct formation could be occurring via a different chemical process. However, the observation of an adduct supports that the formation of disulfide bonds between AC2P20 and other thiol-containing molecules could be occurring in Mtb (Figure 5B).

**Figure 5.**
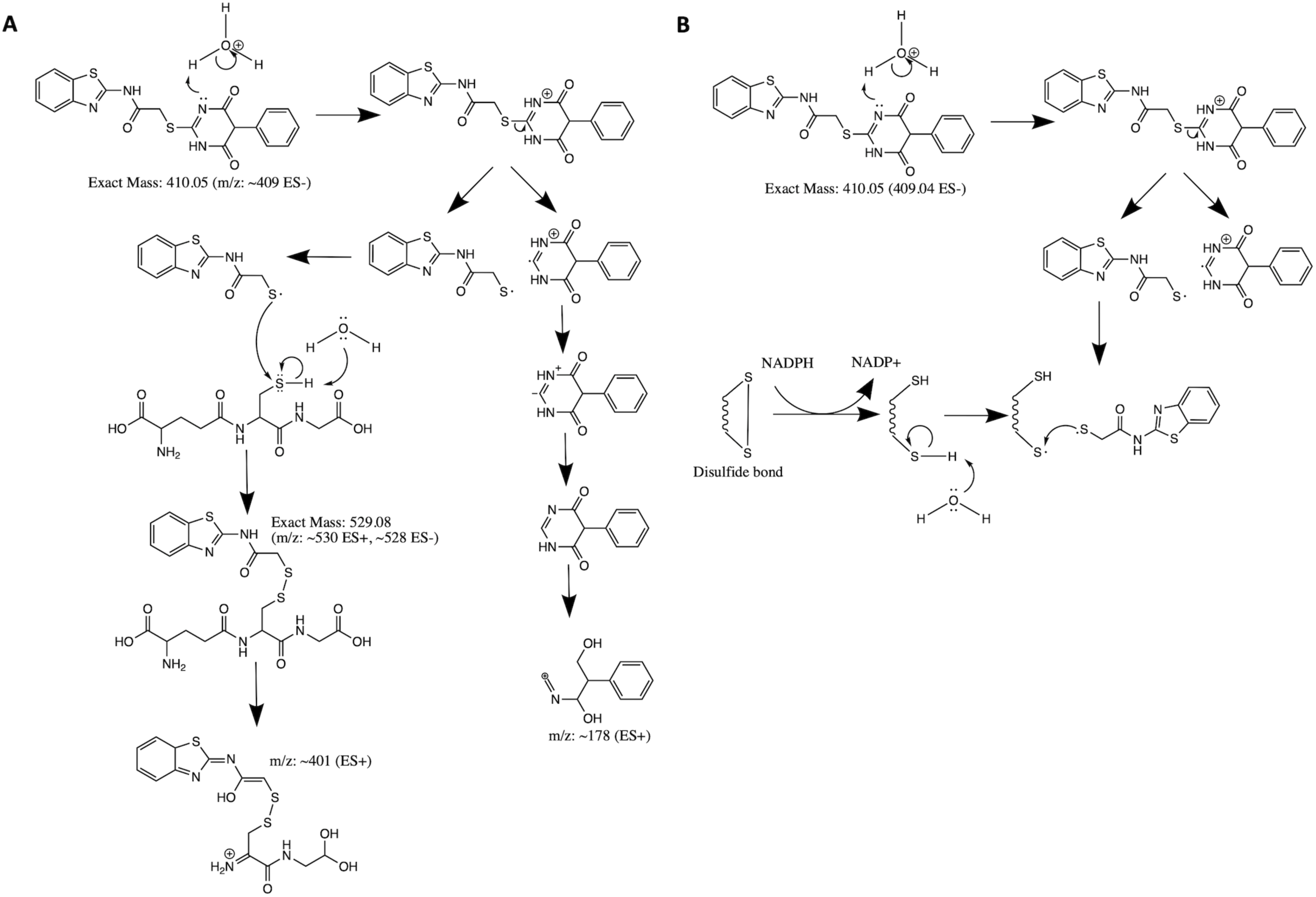
Proposed mechanism for AC2P20 adduct formation. A) Proposed reaction mechanism for the formation of a disulfide bond between AC2P20 and GSH at pH 5.7. B) Proposed stable covalent bond formation between AC2P20 and free thiols in Mtb during redox cycling.

Although, both AC2P20 and AC2P36 function by depleting free thiols, the two scaffolds are distinctly different and engage glutathione (GSH) in different ways. AC2P36 is itself an electrophile, by virtue of the reactive methanesulfonyl moiety on the pyrimidine. GSH can add directly to AC2P36 on the pyrimidine, followed by elimination of the excellent leaving group methanesulfinic acid[29]. On the other hand, AC2P20 is not itself reactive to GSH in an analogous fashion, as evidenced by a lack of MS ion for a direct adduct of GSH to the pyrimidine dione moiety. Instead, AC2P20 has to get hydrolyzed to the free thiol, after which it forms a disulfide with GSH. Therefore, AC2P20 and AC2P26 have different chemical mechanisms of action, and the GSH adducts are chemically distinct (*e.g.* disulfide vs thiopyrimidine).

Redox homeostasis represents a potentially important Mtb vulnerability at acidic pH. Mtb experiences reductive stress during hypoxia and at acidic pH [10]. Genes important for mitigating redox stress are shown to be directly influenced by acid stress; therefore, disruption of redox homeostasis results in the loss of Mtb protection against acid stress [10]. Furthermore, direct perturbations to either redox-homeostasis or pH-homeostasis results in decreased drug tolerance and enhanced Mtb killing [50]. Indeed, chloroquine has recently been shown to kill Mtb *in vivo* by targeting redox homeostasis [50] and auranofin also shows promising antimycobacterial activity [23, 51]. Furthermore, agents targeting respiration may similarly have activity by promoting redox imbalance. Thus, targeting redox-homeostasis represents an important new approach to treating TB. Like AC2P36, we have discovered a second, albeit novel, pH-selective compound (AC2P20) that directly targets free thiols to perturb redox homeostasis. Both AC2P36 and AC2P20 deplete free thiol pools and increase intracellular ROS as part of their killing mechanisms. Interestingly, AC2P20 depletes less free thiols than AC2P36, but has a greater increase in intracellular ROS. This suggests that although both appear to target Mtb free thiols, there are differences in their mechanisms. One hypothesis is that release of the phenyl-dioxopyrimidine group could also be targeting a secondary unknown Mtb physiology, possibly explaining the higher ROS increase that is observed compared to AC2P36 (Figure 4B). Both compounds also form adducts with the low molecular weight thiol, GSH; however, there are major chemical scaffold differences. AC2P36 captures thiols with the release of methylsulfinate while AC2P20 is cleaved to generate benzothioazole-mercaptoacetamide, which then goes on to form disulfide bonds. Although AC2P20 and AC2P36 compounds are structurally unique and have distinct mechanisms-of-action, they do exhibit similar physiological effects on Mtb, supporting the conclusion that thiol redox homeostasis is specifically vulnerable to inhibition at acidic pH.

Several studies in Mtb show a link between low pH- and oxidative stress responses [7,9,29,50,52]. At acidic pH *in vitro*, Mtb exhibits a more reduced cytoplasm and a shift from glycolysis to fatty acid synthesis [9]. This metabolic remodeling is thought to occur in order to generate more oxidized cofactors to mitigate reductive stress. However, a more reduced cytoplasm in Mtb may also play a role in protecting Mtb against oxidative stress. A recent study comparing the RNAseq profiles of reduced MSH redox potential (*E_MSH_*-reduced), intraphagosmal Mtb, and pH stress supports this claim and shows that *E_MSH_*-reduced transcriptome has significant overlap with the pH-regulon[50]. When we compare the *E_MSH_*-reduced, intraphagosomal Mtb, and pH stress regulons with AC2P20 and AC2P36 transcriptional profiles, we again see overlap in redox sensitive genes (i.e. *katG*, *trxB2*, and *whiB3*) which are important for protection against oxidative stress.

While both AC2P20 and AC2P36 share these similar gene induction characteristics, there are differences in specific thiol-related genes. For example, methionine synthesis (i.e. *metK*, *metA*, *metC*) appears modulated by AC2P36 treatment, but induction of these genes is absent in AC2P20 transcriptional data. Likewise, AC2P20 strongly induces sulfate reduction via APS (*cysH*, *nirA*), however, these genes are not modulated by AC2P36. These differences may reflect differences in how these compounds sequester free thiols and which free thiols in particular are being modified. While mycothiol is the most abundant free thiol in Mtb (present in millimolar amounts) [53], it is plausible AC2P20 targets other low molecular weight thiols such as ergothioneine (ERG) [32] or gamma-glutamylcysteine (GGC) [54]. Our mass spectrometry data also supports AC2P20 may be generally targeting free thiols, forming adducts with both GSH and NAC, which would indicate that 1) AC2P20 can target a thiol group in general, and 2) it can directly target a cysteine derivative. Further profiling experiments would need to be undertaken to determine in which molecular contexts AC2P20 targets free thiols and indeed, other related molecules are being developed as tools for cysteine-reactive profiling [45, 46].

## Conclusions

The discovery of two independent molecules selectively killing Mtb at acidic pH by depleting free thiols provides further support for our hypothesis that Mtb is highly sensitive to thiol homeostasis stress at acidic pH and this pathway is a valuable new target for TB drug discovery. AC2P20 or AC2P36 in their present state, will not likely make useful drugs, as they could react with host thiols and thus be neutralized prior to reaching Mtb or could be cytotoxic. However, they independently point the way to further efforts to target this pathway. Indeed, the thioredoxin reductase inhibitor auranofin is in early clinical trials to treat TB and acts, as shown here, similarly functions by depleting free thiols, by a distinct, indirect mechanism. Several groups are pursuing compounds that have enhanced killing at acidic pH but have mostly focused on bacterial pH-homeostasis [16–18]. This new work further validates targeting thiol homeostasis as an alternative target to kill Mtb at acidic pH. Other chemotypes, such as auranofin, that do so indirectly are likely the most promising route. However, it could be possible to develop the compounds related to AC2P20 or AC2P36 into prodrugs that are activated by a Mtb specific enzyme, thus releasing the thiol-reactive warhead selectively inside the bacterial cell. Notably, for both AC2P20 and AC2P36 we could not isolate resistant mutants. This is consistent with the compounds having a broad target (free thiols) and not a specific protein, where resistant mutants could be selected. Therefore, it is possible that should a compound targeting free thiols be developed, the evolution of resistance may be slower as compared to a traditional antibiotic.

In conclusion, our findings have uncovered a novel thiol-targeting chemical probe, AC2P20. AC2P20, in combination with AC2P36, can be classified as a new class of compounds that render Mtb especially sensitive to changes in thiol homeostasis at acidic pH. Further experiments to examine the mechanism of this sensitivity can be undertaken using AC2P20 or AC2P36 as chemical probes. For example, using TN-seq, identification of mutants that become sensitive to AC2P20 and AC2P36 at a neutral pH or have enhanced sensitivity at acidic pH, may reveal key functional pathways required for maintaining thiol-homeostasis.

## Acknowledgements

We thank Christopher Colvin and Javiera Ortiz for technical assistance on the high throughput screening and cytoplasmic pH assays, respectively. Research on this project was supported grants from the NIH-NIAID to RBA (U54AI057153, R01AI116605, R21AI105867) and AgBioResearch.

## Author Contributions

S.J.D., G.B.C., and R.B.A. conceived the project. S.J.D performed the time-dependent and concentration-dependent killing assays, RNAseq analysis, mass spectrometry, free thiol assay, and ROS assay. G.B.C. conducted the initial characterization studies including Mtb and *M. smegmatis* EC_50_ assays and the RNAseq experiment. M.W.W. and S.D.L. contributed to mass spectrometry analysis and proposed mechanism. S.J.D. and R.B.A. wrote the manuscript.

## Supplemental Figures

## Supplemental Table

**Table S1:** Analyzed RNA-seq transcriptional profiling data. S1A. Classification of differentially regulated genes. S1B. Genes significantly induced by AC2P20. S1C. Genes significantly repressed by AC2P20. S1D. Complete list of unfiltered gene expression data in response to AC2P20 treatment.

---

**Supplemental Figure 1.**
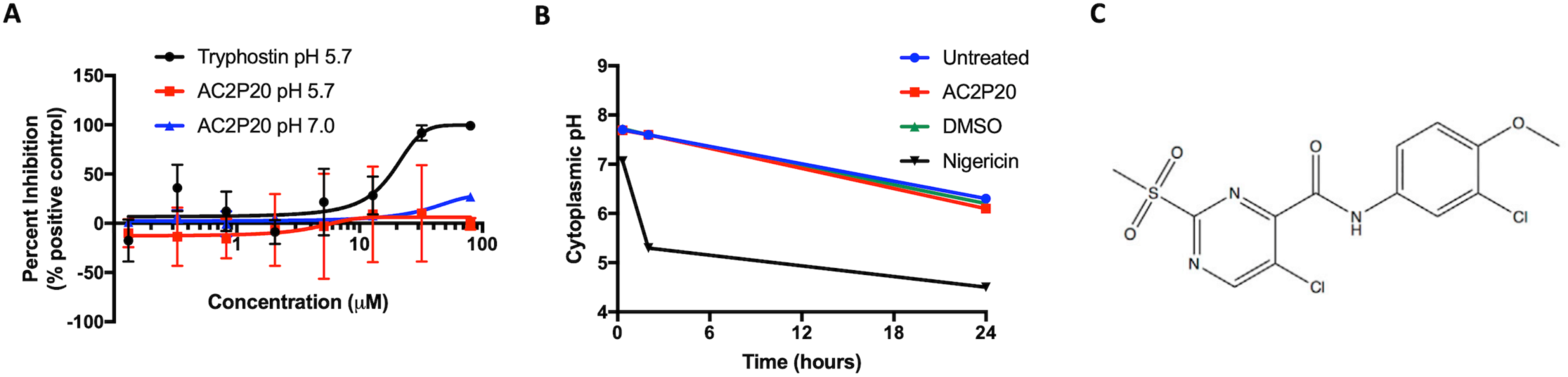
**S1A.** Dose-response curve for AC2P20 inhibition of *M. smegmatis* GFP fluorescence. **S1B.** AC2P20 does not modulate Mtb cytoplasmic pH at pH 5.7. DMSO and Nigericin served as negative and positive controls, respectively. **S1C.** The chemical structure of AC2P36 (5-chloro-N-(3-chloro-4-methoxyphenyl)-2-methylsulfonylpyrimidine-4-carboxamide) [29].

**Supplemental Figure 2.**
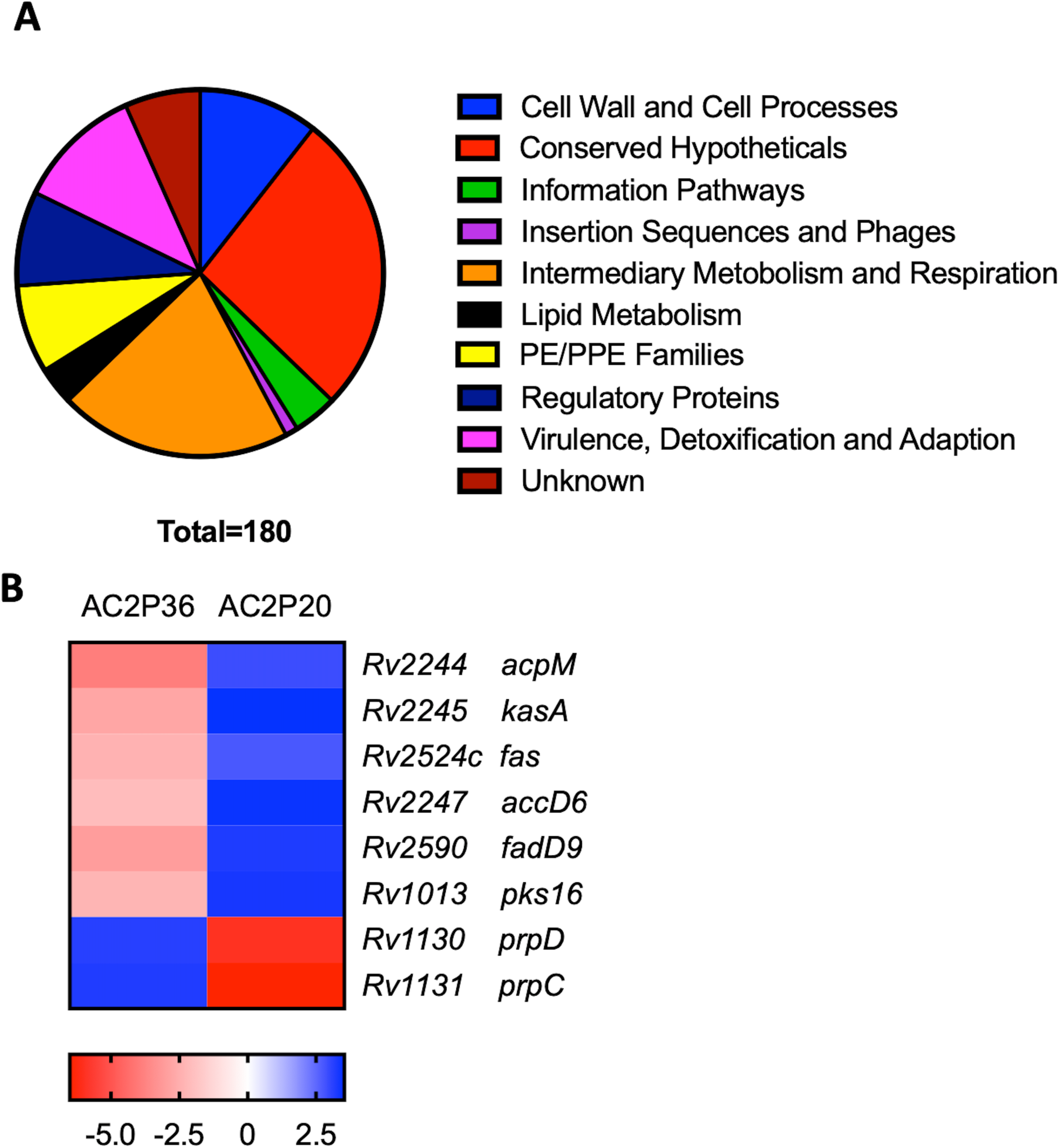
**S2A.** A pie chart depicting the functional classification breakdown of significantly induced genes (>2-fold, q < 0.05) following the analysis of AC2P36-treated Mtb RNA-seq profile. **S2B.** Heatmap comparing the contrast between 8 differentially-regulated genes (between AC2P20 and AC2P36 at pH 5.7) that are involved in lipid metabolism and central metabolism. Genes were annotated with the H37Rv genome.

**Supplemental Figure 3.**
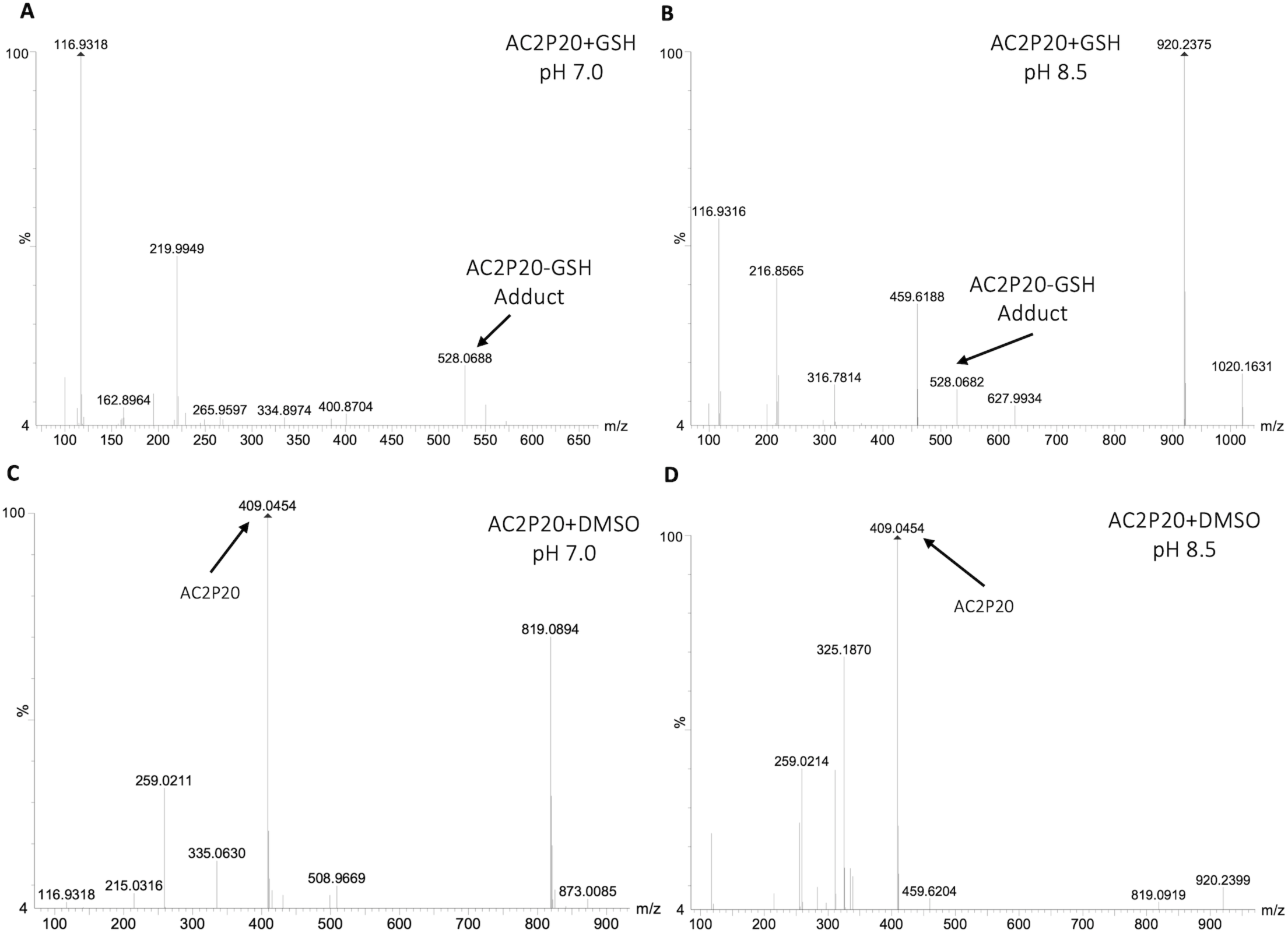
**S3A.** Mass spectrometry data showing adduct formation between AC2P20 and GSH at pH 7.0. Spectra were analyzed in negative ESI mode. **S3B.** Mass spectrometry data showing adduct formation between AC2P20 and GSH at pH 8.5. Spectra were analyzed in negative ESI mode. **S3C.** AC2P20 incubated with DMSO does not fragment in the absence of GSH at pH 7.0. Spectra were analyzed in negative ESI mode. **S3D.** AC2P20 incubated with DMSO does not fragment in the absence of GSH at pH 8.5. Spectra were analyzed in negative ESI mode.

**Supplemental Figure 4.**
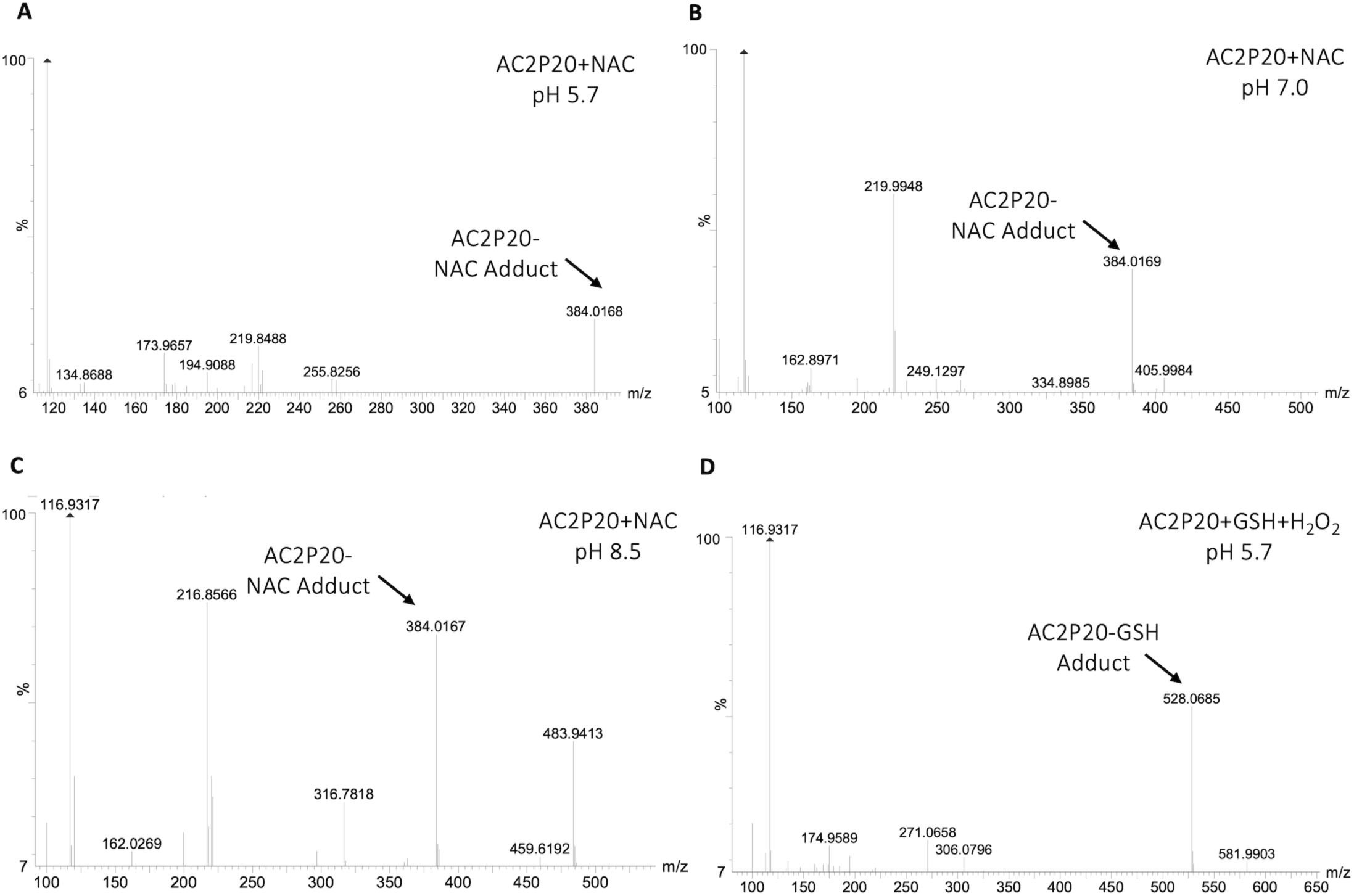
**S4A.** Mass spectrometry data showing adduct formation between AC2P20 and N-acetylcysteine at pH 5.7. Spectra were analyzed in negative ESI mode. **S4B.** Mass spectrometry data showing adduct formation between AC2P20 and N-acetylcysteine at pH 7.0. Spectra were analyzed in negative ESI mode. 20 +NAC 7.0 **S4C.** Mass spectrometry data showing adduct formation between AC2P20 and N-acetylcysteine at pH 8.5. Spectra were analyzed in negative ESI mode. **S4D.** AC2P20 is still able to form an adduct with GSH in the presence of the oxidant, H_2_O_2_. Spectra were analyzed in negative ESI mode.

**Supplemental Figure 5.**
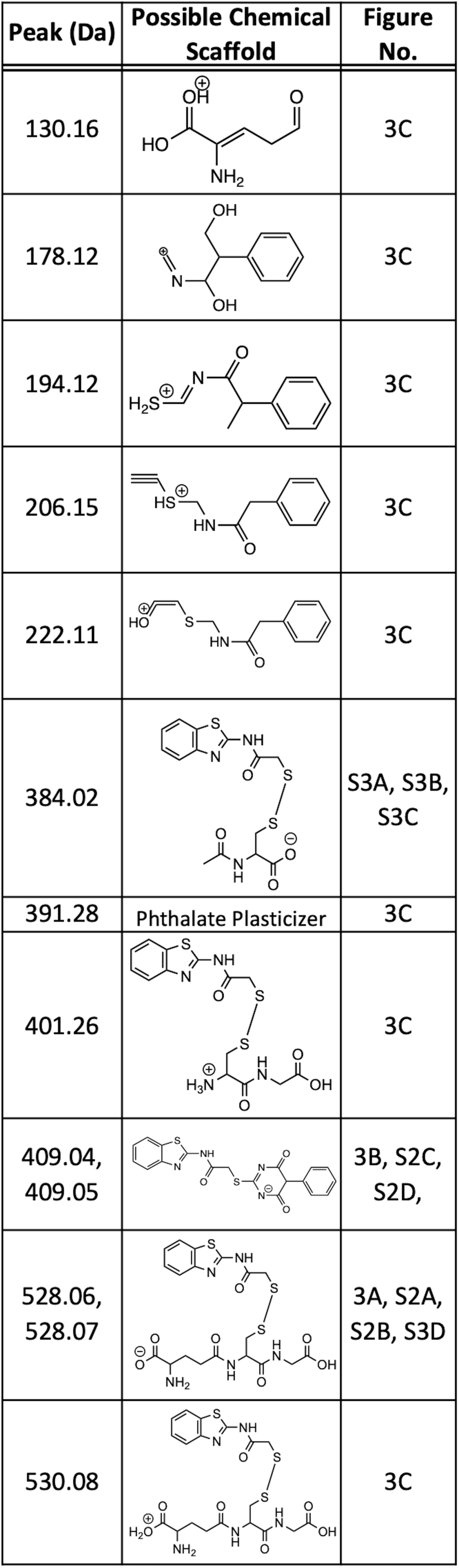
**S5.** A list of labeled mass spectrometry peaks with their corresponding hypothetical chemical scaffolds.

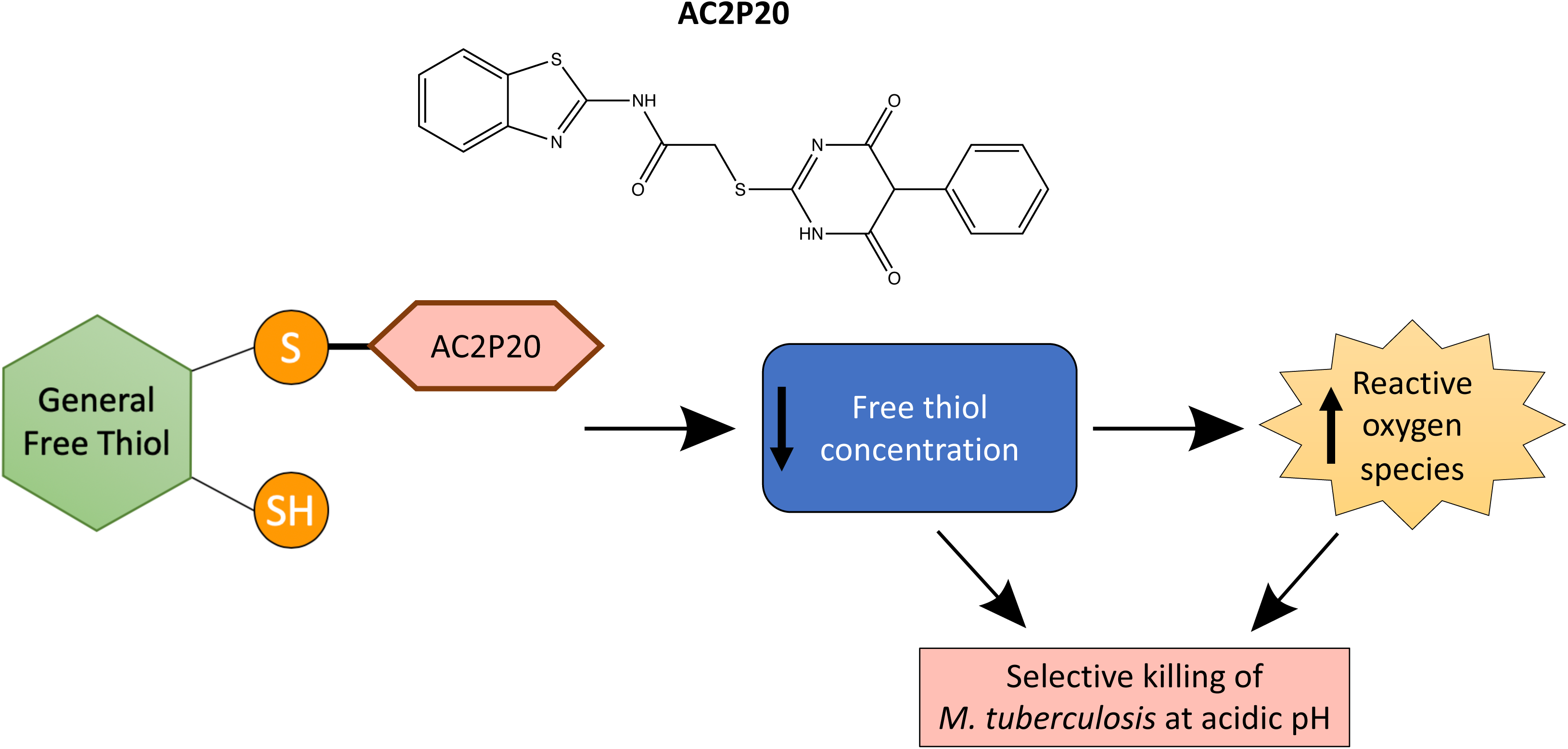

## References

1. Dutta NK, Karakousis PC (2014) Latent tuberculosis infection: myths, models, and molecular mechanisms. Microbiol Mol Biol Rev 78 (3):343–371. doi:10.1128/MMBR.00010-14

2. Rohde KH, Abramovitch RB, Russell DG (2007) Mycobacterium tuberculosis invasion of macrophages: linking bacterial gene expression to environmental cues. Cell Host Microbe 2 (5):352–364. doi:10.1016/j.chom.2007.09.006

3. Baker JJ, Dechow SJ, Abramovitch RB (2019) Acid Fasting: Modulation of Mycobacterium tuberculosis Metabolism at Acidic pH. Trends Microbiol 27 (11):942–953. doi:10.1016/j.tim.2019.06.005

4. MacMicking JD, Taylor GA, McKinney JD (2003) Immune control of tuberculosis by IFN-gamma-inducible LRG-47. Science 302 (5645):654–659. doi:10.1126/science.1088063

5. Vandal OH, Nathan CF, Ehrt S (2009) Acid resistance in Mycobacterium tuberculosis. J Bacteriol 191 (15):4714–4721. doi:10.1128/JB.00305-09

6. Vandal OH, Pierini LM, Schnappinger D, Nathan CF, Ehrt S (2008) A membrane protein preserves intrabacterial pH in intraphagosomal Mycobacterium tuberculosis. Nat Med 14 (8):849–854. doi:10.1038/nm.1795

7. Vandal OH, Roberts JA, Odaira T, Schnappinger D, Nathan CF, Ehrt S (2009) Acid-susceptible mutants of Mycobacterium tuberculosis share hypersusceptibility to cell wall and oxidative stress and to the host environment. J Bacteriol 191 (2):625–631. doi:10.1128/JB.00932-08

8. Song H, Huff J, Janik K, Walter K, Keller C, Ehlers S, Bossmann SH, Niederweis M (2011) Expression of the ompATb operon accelerates ammonia secretion and adaptation of Mycobacterium tuberculosis to acidic environments. Mol Microbiol 80 (4):900–918. doi:10.1111/j.1365-2958.2011.07619.x

9. Baker JJ, Johnson BK, Abramovitch RB (2014) Slow growth of Mycobacterium tuberculosis at acidic pH is regulated by phoPR and host-associated carbon sources. Mol Microbiol 94 (1):56–69. doi:10.1111/mmi.12688

10. Mehta M, Rajmani RS, Singh A (2016) Mycobacterium tuberculosis WhiB3 Responds to Vacuolar pH-induced Changes in Mycothiol Redox Potential to Modulate Phagosomal Maturation and Virulence. J Biol Chem 291 (6):2888–2903. doi:10.1074/jbc.M115.684597

11. Farhana A, Guidry L, Srivastava A, Singh A, Hondalus MK, Steyn AJ (2010) Reductive stress in microbes: implications for understanding Mycobacterium tuberculosis disease and persistence. Adv Microb Physiol 57 (C):43–117. doi:10.1016/B978-0-12-381045-8.00002-3

12. Abramovitch RB, Rohde KH, Hsu FF, Russell DG (2011) aprABC: a Mycobacterium tuberculosis complex-specific locus that modulates pH-driven adaptation to the macrophage phagosome. Mol Microbiol 80 (3):678–694. doi:10.1111/j.1365-2958.2011.07601.x

13. Singh A, Guidry L, Narasimhulu KV, Mai D, Trombley J, Redding KE, Giles GI, Lancaster JR, Jr., Steyn AJ (2007) Mycobacterium tuberculosis WhiB3 responds to O2 and nitric oxide via its [4Fe-4S] cluster and is essential for nutrient starvation survival. Proc Natl Acad Sci U S A 104 (28):11562–11567. doi:10.1073/pnas.0700490104

14. Singh A, Crossman DK, Mai D, Guidry L, Voskuil MI, Renfrow MB, Steyn AJ (2009) Mycobacterium tuberculosis WhiB3 maintains redox homeostasis by regulating virulence lipid anabolism to modulate macrophage response. PLoS Pathog 5 (8):e1000545. doi:10.1371/journal.ppat.1000545

15. Baker JJ, Abramovitch RB (2018) Genetic and metabolic regulation of Mycobacterium tuberculosis acid growth arrest. Sci Rep 8 (1):4168. doi:10.1038/s41598-018-22343-4

16. Zhao N, Darby CM, Small J, Bachovchin DA, Jiang X, Burns-Huang KE, Botella H, Ehrt S, Boger DL, Anderson ED, Cravatt BF, Speers AE, Fernandez-Vega V, Hodder PS, Eberhart C, Rosen H, Spicer TP, Nathan CF (2015) Target-based screen against a periplasmic serine protease that regulates intrabacterial pH homeostasis in Mycobacterium tuberculosis. ACS Chem Biol 10 (2):364–371. doi:10.1021/cb500746z

17. Darby CM, Ingolfsson HI, Jiang X, Shen C, Sun M, Zhao N, Burns K, Liu G, Ehrt S, Warren JD, Andersen OS, Brickner SJ, Nathan C (2013) Whole cell screen for inhibitors of pH homeostasis in Mycobacterium tuberculosis. PLoS One 8 (7):e68942. doi:10.1371/journal.pone.0068942

18. Early J, Ollinger J, Darby C, Alling T, Mullen S, Casey A, Gold B, Ochoada J, Wiernicki T, Masquelin T, Nathan C, Hipskind PA, Parish T (2019) Identification of Compounds with pH-Dependent Bactericidal Activity against Mycobacterium tuberculosis. ACS Infect Dis 5 (2):272–280. doi:10.1021/acsinfecdis.8b00256

19. Reichlen MJ, Leistikow RL, Scobey MS, Born SEM, Voskuil MI (2017) Anaerobic Mycobacterium tuberculosis Cell Death Stems from Intracellular Acidification Mitigated by the DosR Regulon. J Bacteriol 199 (23). doi:10.1128/JB.00320-17

20. Tan MP, Sequeira P, Lin WW, Phong WY, Cliff P, Ng SH, Lee BH, Camacho L, Schnappinger D, Ehrt S, Dick T, Pethe K, Alonso S (2010) Nitrate respiration protects hypoxic Mycobacterium tuberculosis against acid- and reactive nitrogen species stresses. PLoS One 5 (10):e13356. doi:10.1371/journal.pone.0013356

21. Hards K, McMillan DGG, Schurig-Briccio LA, Gennis RB, Lill H, Bald D, Cook GM (2018) Ionophoric effects of the antitubercular drug bedaquiline. Proc Natl Acad Sci U S A 115 (28):7326–7331. doi:10.1073/pnas.1803723115

22. Flentie K, Harrison GA, Tukenmez H, Livny J, Good JAD, Sarkar S, Zhu DX, Kinsella RL, Weiss LA, Solomon SD, Schene ME, Hansen MR, Cairns AG, Kulen M, Wixe T, Lindgren AEG, Chorell E, Bengtsson C, Krishnan KS, Hultgren SJ, Larsson C, Almqvist F, Stallings CL (2019) Chemical disarming of isoniazid resistance in Mycobacterium tuberculosis. Proc Natl Acad Sci U S A 116 (21):10510–10517. doi:10.1073/pnas.1818009116

23. Harbut MB, Vilcheze C, Luo X, Hensler ME, Guo H, Yang B, Chatterjee AK, Nizet V, Jacobs WR, Jr., Schultz PG, Wang F (2015) Auranofin exerts broad-spectrum bactericidal activities by targeting thiol-redox homeostasis. Proc Natl Acad Sci U S A 112 (14):4453–4458. doi:10.1073/pnas.1504022112

24. Lin K, O’Brien KM, Trujillo C, Wang R, Wallach JB, Schnappinger D, Ehrt S (2016) Mycobacterium tuberculosis Thioredoxin Reductase Is Essential for Thiol Redox Homeostasis but Plays a Minor Role in Antioxidant Defense. PLoS Pathog 12 (6):e1005675. doi:10.1371/journal.ppat.1005675

25. Perez E, Samper S, Bordas Y, Guilhot C, Gicquel B, Martin C (2001) An essential role for phoP in Mycobacterium tuberculosis virulence. Mol Microbiol 41 (1):179–187. doi:10.1046/j.1365-2958.2001.02500.x

26. Gonzalo-Asensio J, Mostowy S, Harders-Westerveen J, Huygen K, Hernández-Pando R, Thole J, Behr M, Gicquel B, Martín C (2008) PhoP: A Missing Piece in the Intricate Puzzle of Mycobacterium tuberculosis Virulence. PLOS ONE 3 (10):e3496. doi:10.1371/journal.pone.0003496

27. Walters SB, Dubnau E, Kolesnikova I, Laval F, Daffe M, Smith I (2006) The Mycobacterium tuberculosis PhoPR two-component system regulates genes essential for virulence and complex lipid biosynthesis. Mol Microbiol 60 (2):312–330. doi:10.1111/j.1365-2958.2006.05102.x

28. Johnson BK, Colvin CJ, Needle DB, Mba Medie F, Champion PA, Abramovitch RB (2015) The Carbonic Anhydrase Inhibitor Ethoxzolamide Inhibits the Mycobacterium tuberculosis PhoPR Regulon and Esx-1 Secretion and Attenuates Virulence. Antimicrob Agents Chemother 59 (8):4436–4445. doi:10.1128/AAC.00719-15

29. Coulson GB, Johnson BK, Zheng H, Colvin CJ, Fillinger RJ, Haiderer ER, Hammer ND, Abramovitch RB (2017) Targeting Mycobacterium tuberculosis Sensitivity to Thiol Stress at Acidic pH Kills the Bacterium and Potentiates Antibiotics. Cell Chem Biol 24 (8):993–1004 e1004. doi:10.1016/j.chembiol.2017.06.018

30. Johnson BK, Scholz MB, Teal TK, Abramovitch RB (2016) SPARTA: Simple Program for Automated reference-based bacterial RNA-seq Transcriptome Analysis. BMC Bioinformatics 17:66. doi:10.1186/s12859-016-0923-y

31. Purdy GE, Niederweis M, Russell DG (2009) Decreased outer membrane permeability protects mycobacteria from killing by ubiquitin-derived peptides. Mol Microbiol 73 (5):844–857. doi:10.1111/j.1365-2958.2009.06801.x

32. Saini V, Cumming BM, Guidry L, Lamprecht DA, Adamson JH, Reddy VP, Chinta KC, Mazorodze JH, Glasgow JN, Richard-Greenblatt M, Gomez-Velasco A, Bach H, Av-Gay Y, Eoh H, Rhee K, Steyn AJC (2016) Ergothioneine Maintains Redox and Bioenergetic Homeostasis Essential for Drug Susceptibility and Virulence of Mycobacterium tuberculosis. Cell Rep 14 (3):572–585. doi:10.1016/j.celrep.2015.12.056

33. Williams JT, Haiderer ER, Coulson GB, Conner KN, Ellsworth E, Chen C, Alvarez-Cabrera N, Li W, Jackson M, Dick T, Abramovitch RB (2019) Identification of New MmpL3 Inhibitors by Untargeted and Targeted Mutant Screens Defines MmpL3 Domains with Differential Resistance. Antimicrob Agents Chemother 63 (10). doi:10.1128/AAC.00547-19

34. Zheng H, Colvin CJ, Johnson BK, Kirchhoff PD, Wilson M, Jorgensen-Muga K, Larsen SD, Abramovitch RB (2017) Inhibitors of Mycobacterium tuberculosis DosRST signaling and persistence. Nat Chem Biol 13 (2):218–225. doi:10.1038/nchembio.2259

35. Kapopoulou A, Lew JM, Cole ST (2011) The MycoBrowser portal: a comprehensive and manually annotated resource for mycobacterial genomes. Tuberculosis (Edinb) 91 (1):8–13. doi:10.1016/j.tube.2010.09.006

36. Voskuil MI, Bartek IL, Visconti K, Schoolnik GK (2011) The response of mycobacterium tuberculosis to reactive oxygen and nitrogen species. Front Microbiol 2:105. doi:10.3389/fmicb.2011.00105

37. Manganelli R, Voskuil MI, Schoolnik GK, Dubnau E, Gomez M, Smith I (2002) Role of the extracytoplasmic-function sigma factor sigma(H) in Mycobacterium tuberculosis global gene expression. Mol Microbiol 45 (2):365–374. doi:10.1046/j.1365-2958.2002.03005.x

38. Raman S, Song T, Puyang X, Bardarov S, Jacobs WR, Jr., Husson RN (2001) The alternative sigma factor SigH regulates major components of oxidative and heat stress responses in Mycobacterium tuberculosis. J Bacteriol 183 (20):6119–6125. doi:10.1128/JB.183.20.6119-6125.2001

39. Voss M, Nimtz M, Leimkuhler S (2011) Elucidation of the dual role of Mycobacterial MoeZR in molybdenum cofactor biosynthesis and cysteine biosynthesis. PLoS One 6 (11):e28170. doi:10.1371/journal.pone.0028170

40. Warrier T, Kapilashrami K, Argyrou A, Ioerger TR, Little D, Murphy KC, Nandakumar M, Park S, Gold B, Mi J, Zhang T, Meiler E, Rees M, Somersan-Karakaya S, Porras-De Francisco E, Martinez-Hoyos M, Burns-Huang K, Roberts J, Ling Y, Rhee KY, Mendoza-Losana A, Luo M, Nathan CF (2016) N-methylation of a bactericidal compound as a resistance mechanism in Mycobacterium tuberculosis. Proc Natl Acad Sci U S A 113 (31):E4523–4530. doi:10.1073/pnas.1606590113

41. Sun Z, Cheng SJ, Zhang H, Zhang Y (2001) Salicylate uniquely induces a 27-kDa protein in tubercle bacillus. FEMS Microbiol Lett 203 (2):211–216. doi:10.1111/j.1574-6968.2001.tb10843.x

42. Cole ST, Brosch R, Parkhill J, Garnier T, Churcher C, Harris D, Gordon SV, Eiglmeier K, Gas S, Barry CE, 3rd, Tekaia F, Badcock K, Basham D, Brown D, Chillingworth T, Connor R, Davies R, Devlin K, Feltwell T, Gentles S, Hamlin N, Holroyd S, Hornsby T, Jagels K, Krogh A, McLean J, Moule S, Murphy L, Oliver K, Osborne J, Quail MA, Rajandream MA, Rogers J, Rutter S, Seeger K, Skelton J, Squares R, Squares S, Sulston JE, Taylor K, Whitehead S, Barrell BG (1998) Deciphering the biology of Mycobacterium tuberculosis from the complete genome sequence. Nature 393 (6685):537-544. doi:10.1038/31159

43. Garbe TR (2004) Co-induction of methyltransferase Rv0560c by naphthoquinones and fibric acids suggests attenuation of isoprenoid quinone action in Mycobacterium tuberculosis. Can J Microbiol 50 (10):771–778. doi:10.1139/w04-067

44. Camus JC, Pryor MJ, Medigue C, Cole ST (2002) Re-annotation of the genome sequence of Mycobacterium tuberculosis H37Rv. Microbiology 148 (Pt 10):2967–2973. doi:10.1099/00221287-148-10-2967

45. Motiwala HF, Kuo YH, Stinger BL, Palfey BA, Martin BR (2020) Tunable Heteroaromatic Sulfones Enhance in-Cell Cysteine Profiling. J Am Chem Soc 142 (4):1801–1810. doi:10.1021/jacs.9b08831

46. Zambaldo C, Vinogradova EV, Qi X, Iaconelli J, Suciu RM, Koh M, Senkane K, Chadwick SR, Sanchez BB, Chen JS, Chatterjee AK, Liu P, Schultz PG, Cravatt BF, Bollong MJ (2020) 2-Sulfonylpyridines as Tunable, Cysteine-Reactive Electrophiles. J Am Chem Soc 142 (19):8972–8979. doi:10.1021/jacs.0c02721

47. Murphy CM, Fenselau C, Gutierrez PL (1992) Fragmentation characteristic of glutathione conjugates activated by high-energy collisions. J Am Soc Mass Spectrom 3 (8):815–822. doi:10.1016/1044-0305(92)80004-5

48. Fine Z, Wood TD (2013) Formation of Mercury(II)-Glutathione Conjugates Examined Using High Mass Accuracy Mass Spectrometry. Int J Anal Mass Spectrom Cromatogr 1 (2):90–94. doi:10.4236/ijamsc.2013.12011

49. Abedinzadeh Z, Gardes-Albert M, Ferradini C (1989) Kinetic study of the oxidation mechanism of glutathione by hydrogen peroxide in neutral aqueous medium. Canadian Journal of Chemistry 67:1247–1255

50. Mishra R, Kohli S, Malhotra N, Bandyopadhyay P, Mehta M, Munshi M, Adiga V, Ahuja VK, Shandil RK, Rajmani RS, Seshasayee ASN, Singh A (2019) Targeting redox heterogeneity to counteract drug tolerance in replicating Mycobacterium tuberculosis. Sci Transl Med 11 (518). doi:10.1126/scitranslmed.aaw6635

51. Ruth MM, van Rossum M, Koeken V, Pennings LJ, Svensson EM, Ruesen C, Bowles EC, Wertheim HFL, Hoefsloot W, van Ingen J (2019) Auranofin Activity Exposes Thioredoxin Reductase as a Viable Drug Target in Mycobacterium abscessus. Antimicrob Agents Chemother 63 (9). doi:10.1128/AAC.00449-19

52. Small JL, O’Donoghue AJ, Boritsch EC, Tsodikov OV, Knudsen GM, Vandal O, Craik CS, Ehrt S (2013) Substrate specificity of MarP, a periplasmic protease required for resistance to acid and oxidative stress in Mycobacterium tuberculosis. J Biol Chem 288 (18):12489–12499. doi:10.1074/jbc.M113.456541

53. Newton GL, Arnold K, Price MS, Sherrill C, Delcardayre SB, Aharonowitz Y, Cohen G, Davies J, Fahey RC, Davis C (1996) Distribution of thiols in microorganisms: mycothiol is a major thiol in most actinomycetes. J Bacteriol 178 (7):1990–1995. doi:10.1128/jb.178.7.1990-1995.1996

54. Sao Emani C, Williams MJ, Van Helden PD, Taylor MJC, Wiid IJ, Baker B (2018) Gamma-glutamylcysteine protects ergothioneine-deficient Mycobacterium tuberculosis mutants against oxidative and nitrosative stress. Biochem Biophys Res Commun 495 (1):174–178. doi:10.1016/j.bbrc.2017.10.163

